# Test of a conservation intervention highlights temporal variability in hybridization dynamics in *Catostomus* fishes

**DOI:** 10.1101/2024.07.19.604301

**Authors:** Jillian N. Campbell, Zachary E. Hooley-Underwood, Elizabeth G. Mandeville

## Abstract

Non-native species are a leading threat to fish biodiversity. They pose risks to native populations through human-mediated introductions resulting in hybridization events, which could result in demographic or genetic swamping. *Catostomus* fishes in the Upper Colorado River Basin are an example of this. Extensive hybridization occurs between non-native white suckers (*C. commersonii* ) and native flannelmouth and bluehead suckers (*C. latipinnis* and *C. discobolus*). This system provides a suitable model for using genomic analyses to test the efficacy of an intervention to reduce the abundance of non-native species and production of hybrid offspring. This study implemented a Resistance Board Weir (RBW) as a fish barrier across Roubideau Creek, a tributary of the Gunnison River in Colorado (USA), to restrict non-native sucker participation in spawning events. Conducted over four years, the study gathered genomic data from larval fish samples, pre- and post-implementation of the RBW. We used genomic data to determine the efficacy of a RBW at limiting non-native and hybridized sucker larval production. We found no significant effect of the weir on the proportion of white sucker ancestry in larval fish across the four years of the study, which included three years of weir usage when access was successfully controlled for variable amounts of time. Overall, this work provides insight into the efficacy of a resistance board weir as a management tool for non-native suckers, and highlights interannual variability. This work contributes valuable information for policy and fisheries management in Colorado.

## Introduction

Introducing non-native fish species into the range of other fish, whether on purpose or accidentally, can have negative impacts on native populations through predation (Vitule et al. 2006), competition (Zimmerman and Vondracek 2006), disease (Hickley and Chare 2004), and hybridization, or interbreeding between different species (Cross 2000). Hybridization in fish is widespread and occurs in both freshwater and marine species. Historically, fresh-water fish species represent more fish hybridization overall (Lagler et al. 1982). Although some instances of hybridization result from natural causes, anthropogenic influences have also played a role in these hybridization events. For instance, humans hybridize fish species in order to manage sport fishing opportunities and support commercial fisheries. The aquaculture industry also breeds genetically distinct fish together with the goal of producing fish that have desirable traits (Bartley et al. 2000).

One problem when working with fish populations is that they can be hard to monitor, especially non-intrusively, or when populations are of conservation concern (Lacoursìere-Roussel et al. 2016). Hybrid fish pose an additional challenge, where they may favour the phenotype of one parent species, which allows them to remain undetected in a population unless genetic analyses are completed (Wilk and Horth 2016). To manage hybridization successfully for conservation in fish, it is imperative that we continue to examine fish DNA for signs of hybridization, especially in species known to have cryptic hybrids.

*Catostomus* suckers are a genus of freshwater fish known to hybridize extensively (Hubbs and Miller 1953, Clarkson and Minckley 1988, McDonald et al. 2008, Dowling et al. 2016, Mandeville et al. 2015, 2017, Bangs et al. 2018). This family of fishes, Catostomidae, is thought to be the result of a whole genome duplication event approximately 50 million years ago after species hybridized (Ferris 1984). Hybridization between native and non-native *Catostomus* suckers is a major problem in the Upper Colorado River Basin that is of management interest. Conservation of the native flannelmouth and bluehead suckers (*C. latipinnis* and *C. discobolus*) against hybridization with the non-native white suckers (*C. commersonii* ) is a priority for state, federal, and tribal agencies (Hooley-Underwood et al. 2019, Thompson and Hooley-Underwood 2019). Previous genetic work suggests that both demographic and genetic swamping might be concerns for these species (Mandeville et al. 2015, 2017).

Conservation of fish species comes in many different forms, and each effort taken to conserve native fish species must be tailored to the specific needs of that case. In this study, we used a barrier intervention to trap and selectively allow passage of fish, while culling non-native individuals. Often times barriers are seen as damaging in fisheries science (e.g. LeMoine et al. 2020, Jones et al. 2021), but there can also be benefits to using them. For example, weirs have been used to successfully control invasive species such as sea lamprey by limiting access of sea lamprey to suitable spawning and larval habitat (Lucas et al. 2009, Zielinski et al. 2019).

In this study, we tested a resistance board weir (RBW) to understand its efficacy at reducing the presence of non-native white and longnose suckers and their hybrids in spawning tributaries, and in turn reducing non-native sucker ancestry in offspring produced. The RBW comprises a barrier and a trap (Tobin 1994). As adult fish are swimming upstream to reach spawning habitat they are blocked by the weir, which spans the width of the creek. The RBW funnels fish attempting to move upstream into a trap in the middle of the weir. Once in the trap, field technicians identify the fish morphologically and if they are believed to have non-native sucker ancestry they are removed from the creek. Suckers believed to be of flannelmouth and bluehead ancestry only (as well as all other native fishes), are then returned to the creek above the RBW, where they are able to travel upstream to suitable spawning habitat. Effectiveness of the weir was measured by assessing larval fish ancestry post-spawning. We expected that during years where the RBW intervention was active, non-native sucker ancestry would be lower than in years that the weir was inactive. Additionally, random sampling and genetic identification of adult fish trapped at the RBW was used to aaccount for how well fish species and hybrids are identified by field staff at the weir.

Over the span of four years, we evaluated the efficacy of the weir to successfully reduce non-native suckers and their hybrids, under varying numbers of operational weir days. Genomic analysis of larval fish sampled post-spawning was used to evaluate the different parental species contribution to the sampled population of larvae. This genomic data was compared across years to determine relative success of the weir at reducing hybridization between native and non-native *Catostomus* suckers.

## Methods

### Site location and sampling

This study took place within the Gunnison River Basin of Colorado (USA). A RBW was used as a fish barrier to selectively allow only suckers phenotypically identified as native to participate in an annual tributary spawning event. The RBW was installed near the mouth of Roubideau Creek, a tributary of the Gunnison River (Fig. S1). The Roubideau Creek drainage is mostly an intermittent system, flowing during the annual snow-melt induced runoff (March - June). Above average annual snowpack, and anthropogenic water use can cause sections of streams in the drainage to have perennial flow, but the vast majority of fish that occur in the drainage are migratory (Hooley-Underwood et al. 2019, Thompson and Hooley-Underwood 2019). Typically, adult fishes immigrate from the Gunnison River during March and April, spawn during April and May, and emigrate during May and June. Larval fish hatch during May and June and passively drift out of the drainage during June and July. Adult and larval sucker sampling by Colorado Parks and Wildlife has identified high levels of use and spawning by suckers in the Roubideau Creek Drainage, compared to other tributaries, and therefore, the creek was identified as a high value conservation target. Additionally, the fact that nearly all sucker use of Roubideau Creek is seasonal is ideal as all spawning fish must pass the site selected for the RBW at the mouth of the creek.

The RBW was designed and constructed by Cramer Fish Sciences (Portland, OR, USA) with bar spacing of 30 mm to prevent upstream passage of all fish *>* 300mm (total length). Colorado Parks and Wildlife (CPW) staff placed the weir annually during the beginning of March before any immigration into Roubideau Creek began. The weir was to be operated until immigration was observed to cease - approximately mid May typically. Operation of the weir included regular observation and cleaning to ensure panels did not submerge due to debris accumulation. Daily, all immigrating fish that were trapped were sorted and processed. Most fish movement occurred during afternoon through early morning. Staff attempted to process fish through the trap as quickly as possible to minimize fish behavioral and health effects of being caged, so most fish processing occurred at night. Processing consisted of morphologically identifying and counting all trapped fish, and then transporting native fishes above the weir or culling all fishes that were presumed to possess non-native ancestry. Additionally, a subset of trapped fishes were genetically sampled.

Sampling of adult and larval *Catostomus* fishes was conducted by CPW. From 2019 to 2022, a subset of spawning adults and their progeny (larval fish) were sampled for genetic material from the weir (adults) and 9 sites upstream of the RBW (larvae). Two sets of adult suckers were sampled to validate phenotypic identification of suckers handled at the RBW. One set (target n=30 per parental species per year) consisted of reference individuals – individuals that were believed to represent genetically pure individuals based on phenotypic characteristics including mouth structure, body shape, scale size and organization, and fin shape and ray counts. The other set (target n=60 per year) consisted of random individuals – individuals that were sampled daily from the total weir catch based on randomly generated numbers that were known only by data recorders, and not by staff who were making identifications. Adult genetic material was collected by taking a roughly 1 cm square clipping of lower caudal fin with flame-sterilized scissors. Fin clips were preserved in 2 ml micro-centrifuge tubes with 80-100% ethanol. Larvae were collected using 250 micron mesh drift nets during night time hours, or by dip-netting with a variety of fine-meshed aquarium nets during daylight hours. Larvae were collected from multiple locations and times to ensure that the whole population was representatively sampled. Larvae were collected whole, and were preserved in the same fashion as the adult fin clips.

2019 represents the year where there was no intervention present, and served as a baseline. During 2019, 240 larval fish were collected after the spawning season. 2020 was the first year with the weir in the creek. During years of intervention, all fish identified phenotypically as being non-native to the system or as a hybrid, involving a non-native fish, were removed from the spawning population. Native suckers were allowed to pass the RBW. The weir was operational for different proportions of the spawning season in different years. Due to the COVID-19 pandemic, in March 2020 the weir was pulled out of Roubideau Creek early, after only 29 days, making 2020 a partially controlled year. However, once spawning was completed 240 larval fish were still collected. In 2021, the weir was in the creek for 78 days, the full length of the spawning season. After spawning had been completed, 355 larval fish were collected. Additionally, 58 random and 100 reference adult fin clips were taken during RBW operation. In 2022, the RBW was installed for an additional year of intervention. Unfortunately, rapid warming in the spring led to fast melting of snow off the mountains and therefore increased stream flows too fast. This caused structural failure of the RBW, as it could not withstand the debris build up and increased flows. Therefore, 2022 represents a partially controlled year with the weir active for 48 days. In total, 472 larval fish were collected after spawning and 43 random and 90 reference adult fin clip samples were collected at the weir. Across all four years a total of 1307 larval fish were sampled and a total of 290 adult fish were sampled, for a grand total of 1597 samples.

### Genomic data preparation

DNA was extracted from all samples using DNeasy Blood & Tissue Kits (Qiagen, Inc.) according to manufacturer’s instructions. For each sample, approximately half a larval fish or half a fin clip was used. Extracted DNA was quantified using a Nanodrop spectrophotometer. The extracted DNA was then used to create highly-multiplexed genomic libraries for high-throughput sequencing. These libraries were prepared following methods from Parchman et al. 2012. This process involved restriction enzyme digest of the prepared samples with EcoRI and MseI, followed by ligation of adaptors, including unique barcodes, to each individual sample. The restriction/ligation product was then amplified using two rounds of PCR (Parchman et al. 2012). The resulting genomic libraries were size selected at a narrow targeted range of 275 base pair fragments (approximately 250–300bp) using a PippinPrep machine (SageScience). The final genomic libraries were then sent to SickKids Toronto for Illumina sequencing on the NovaSeq 6000 at The Centre for Applied Genomics (TCAG). Sequencing yielded large amounts of raw genomic data which was then assessed using bioinformatics.

### Filtering and variant calling

The raw genomic libraries were first demultiplexed using sabre (https://github.com/najoshi/sabre). This process involved matching 8-10 base pair barcodes with samples using text files containing each unique barcode and paired sample ID. The output FASTQ files were then aligned to a reference flannelmouth sucker genome (GenBank GCA 036785435.1; ASM3678543v1, Mandeville et al. 2024) using Burrows-Wheeler alignment (bwa version 0.7.17; Li and Durbin 2010, Li 2013). Prior to alignment, the total number of raw reads was 3.46 *×* 10^9^. After alignment to the reference genome 2.38 *×* 10^9^ reads were retained (approximately 69%). Using the 1643 individual bam files, single nucleotide variants (SNPs) were called using samtools version 1.9 and bcftools version 1.9 (Li et al. 2009, Li 2011). The resulting VCF file was filtered using vcftools version 0.1.16 (Danecek et al. 2011). For the purpose of this study we wanted sites that were polymorphic in multiple populations. Therefore, sites with a minor allele frequency of less than 5% were filtered out. Additionally, sites with more than 2 alleles were removed and only one variable site per contig was randomly selected using the thin parameter (--thin 90). As well, individuals with low-coverage depth (*>*90% missing data) were filtered out. In total, 1,078 individuals and 34,828 SNPs were retained in .vcf format. We further filtered out potentially paralogous loci using vcftools version 0.1.16 and a custom R script (based on the approach in McKinney et al. 2017), as the Catostomidae resulted from a whole-genome duplication. This filtering step removed an additional 5,507 SNPs. Finally, 60 additional individuals were removed from the dataset as these individuals were sampled for another research question. The remaining 29,321 SNPs and 1018 individuals (808 larval, 210 adult) were used for all analyses going forward.

### Ancestry determination using **entropy**

The hierarchical Bayesian model entropy (Gompert et al. 2014, Shastry et al. 2021) was used to make estimates of ancestry for each individual. This model creates estimates of *q* (proportion of ancestry) and *Q* (interspecific ancestry) which are used to distinguish pure, F1, backcrossed, and advanced-generation hybrids. The value of *q* describes the proportion of an individual’s ancestry that comes from each parental species, whereas *Q* describes the proportion of loci in an individual that have ancestry from both parental species. Values of *q* = 1 represents a fish with pure ancestry in one of the parental species. However, other intermediate values of *q* represent a fish with mixed ancestry, i.e. a hybrid. To distinguish clearly what types of hybrids we have, *Q* is important. As *Q* represents interspecific ancestry, we would expect that *Q* = 0 for pure parental species, *Q* = 1 for F1 hybrids, and *Q* = 0.5 for F2 hybrids and first-generation backcrosses.

To begin, the model was run to make estimates of *q* using values of *k* = 1-5. Here, *k* represents the number of genetic clusters, or parental species, that entropy will break the individuals out into. The optimal value of *k* was determined after the model had run using DIC (Deviance Information Criterion) values to assess model fit to these data (Fig. S2). For each value of *k* we ran three replicate runs. We ran MCMC (Markov chain Monte Carlo) chains for 50,000 steps to make posterior distribution estimates. Of those 50,000 steps the first 25,000 were discarded as burn in. After the burn in was complete, every 25th step was retained. Model convergence was checked by plotting MCMC chains for a random subset of individuals (Fig. S3).

Individuals were designated as either one of the parental species or a type of hybrid using a custom script (identification_field_genetic.R). Taking the *q* values, individuals were classified based on the proportion of ancestry belonging to the four different parental species. An individual with *≥*95% ancestry from one parental species was designated as purely one species (ex. pure white sucker). If an individual had *<*95% ancestry from two groups and *<*5% for the other two groups, that individual was designated a hybrid (ex. white*×*flannelmouth hybrid). The remaining individuals, being any fish with *>*5% ancestry in more than two groups, was designated as an “other hybrid” (for example a multispecies hybrid). After individuals were given species estimates using *q* values, entropy was rerun on species pairs (*k* = 2) to get estimates on interspecific ancestry (*Q* ). The same parameters were used, with 50,000 steps of MCMC chains, 25,000 steps of burn in, and every 25th step being retained after the burn in was complete. Due to the very low occurrence of longnose suckers, only three species pairs were evaluated, white *×* flannelmouth, white *×* bluehead, and flannelmouth *×* bluehead. Three replicate runs were also completed on each of the species pairs. We used the relationship between *q* and *Q* to make detailed ancestry estimates of hybrids.

## Results

### Proportions of spawning adults

Data from the resistance board weir operations showed that the majority of the fish arriving in Roubideau creek to spawn were phenotypically identified as parental species. In 2020, a partial control year truncated by the COVID-19 pandemic, 1689 blueheads, 319 flannelmouths, 40 white suckers, 1 longnose sucker, and 329 putative hybrids were identified at the weir. In 2021, when successful operation of the weir spanned the full spawning season, 13676 blueheads, 8702 flannelmouths, 877 white suckers, 17 longnose suckers, and 1347 putative hybrids were identified. In 2022, when high flows forced an early removal of the weir in April, 2644 blueheads, 6388 flannelmouths, 154 white suckers, 0 longnose suckers, and 805 putative hybrids were identified at the weir. This indicates that the relative proportions of native parental species were highly variable across years (blueheads 26–71% of spawners, flannelmouths 13–63%), and that the non-native white suckers and hybrids were a variable but smaller proportion of spawners (1–4% and 5–13% respectively). However, for 2020 and 2022, our data does not cover the entire spawning window, so additional white suckers and hybrids may have entered Roubideau Creek after the weir ceased operation for the season.

### Impact of weir usage was extremely variable across years

Percentage of larval ancestry coming from the different parental species varied substantially across years. In 2019, when the weir was not present, flannelmouth ancestry made up 38.97% of the population; 51.64% was bluehead sucker ancestry, 9.34% white sucker, and 0.05% longnose sucker. In 2020, when the weir was present for 29 days, flannelmouth ancestry represented 28.23% of the population, bluehead ancestry represented 60.01% of the population, while white and longnose sucker represented 11.75% and 0.01%, respectively. In 2021, when the weir was present for the entirety of the season (78 days), flannelmouth ancestry made up 14.84% of the population, 81.34% was bluehead sucker, 3.59% was white sucker, and 0.23% was longnose sucker. Finally, in 2022, when the weir was present for 48 days, flannelmouth sucker ancestry made up 27.90% of the population, bluehead suckers made up 34.02% of the population, 37.80% came from white suckers, and 0.28% from longnose suckers (Fig. 1, Table S1). A Pearson Correlation Coefficient test showed no significance or correlation (*p >* 0.05, R=0) between the amount of time the weir was active in the creek and the mean proportion of white sucker ancestry (Fig. 2).

**Figure 1:**
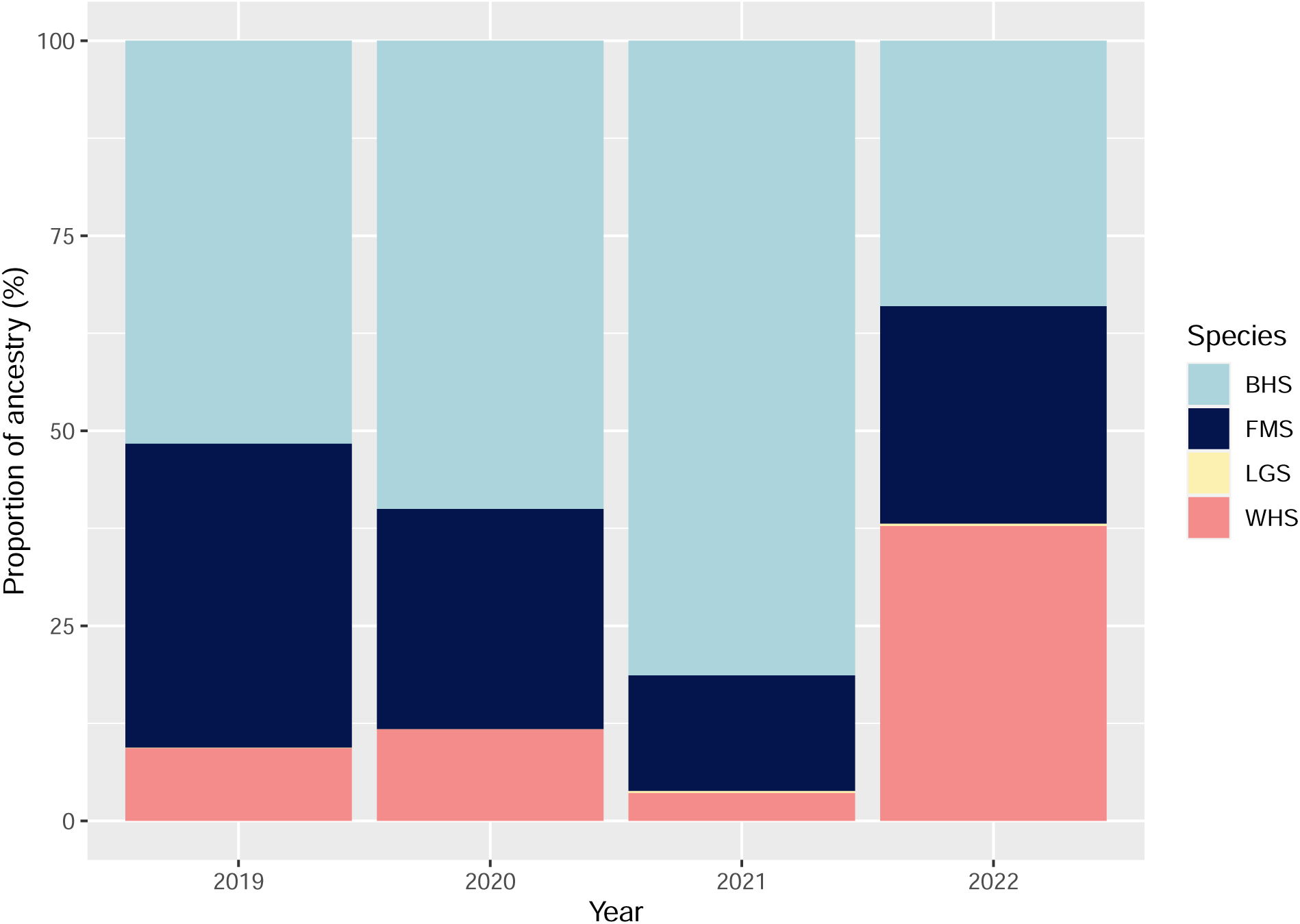
Proportion of ancestry of the sampled larval population across years coming from each of the parental species. Colours represent the four parental species, flannelmouth sucker (FMS), bluehead sucker (BHS), white sucker (WHS), and longnose sucker (LGS).

**Figure 2:**
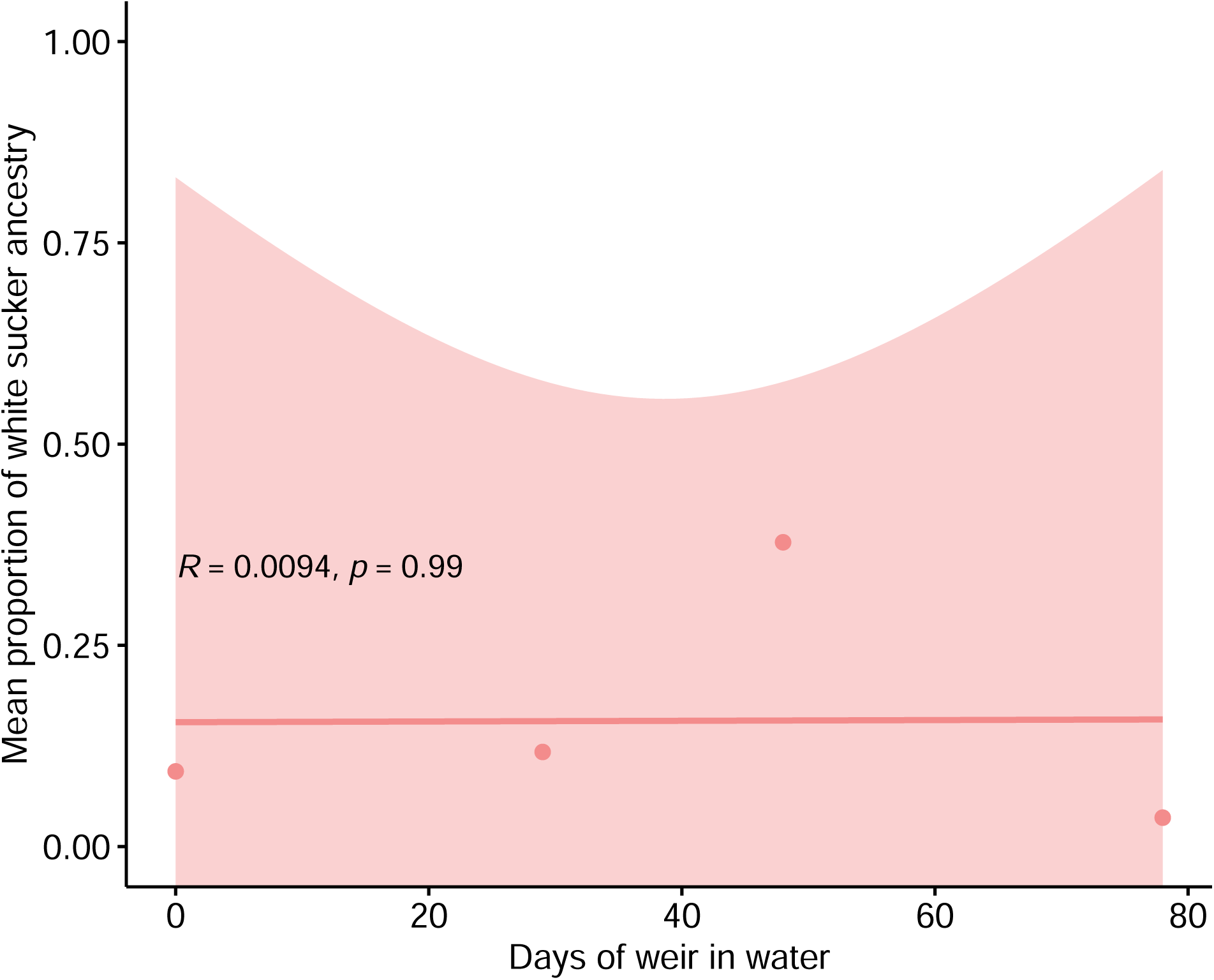
Average proportion of white sucker ancestry in larval fish across varying degrees of weir usage. Individual points represent the percentage of the total larval population with white sucker ancestry during each of the four different amounts of time the weir spent in the creek. The line represents a Pearson Correlation Coefficient test to look for significance between the two variables. CIs (confidence intervals) are represented by the pink shaded area. There is no significance or correlation between weir days and average amount of white sucker ancestry in the population (*p >* 0.05, R=0).

The program entropy allowed for visual representation of larval ancestry data across years (Fig. 3). Ancestry proportions coming from the different species can be seen changing between years. The non-native white sucker was most prominent in 2022 and least in 2021. Additionally, the non-native longnose sucker contributed most ancestry in 2021 and 2022, with virtually no longnose sucker ancestry within the sampled population during 2019 and 2020 (Fig. 1, Table S1).

**Figure 3:**
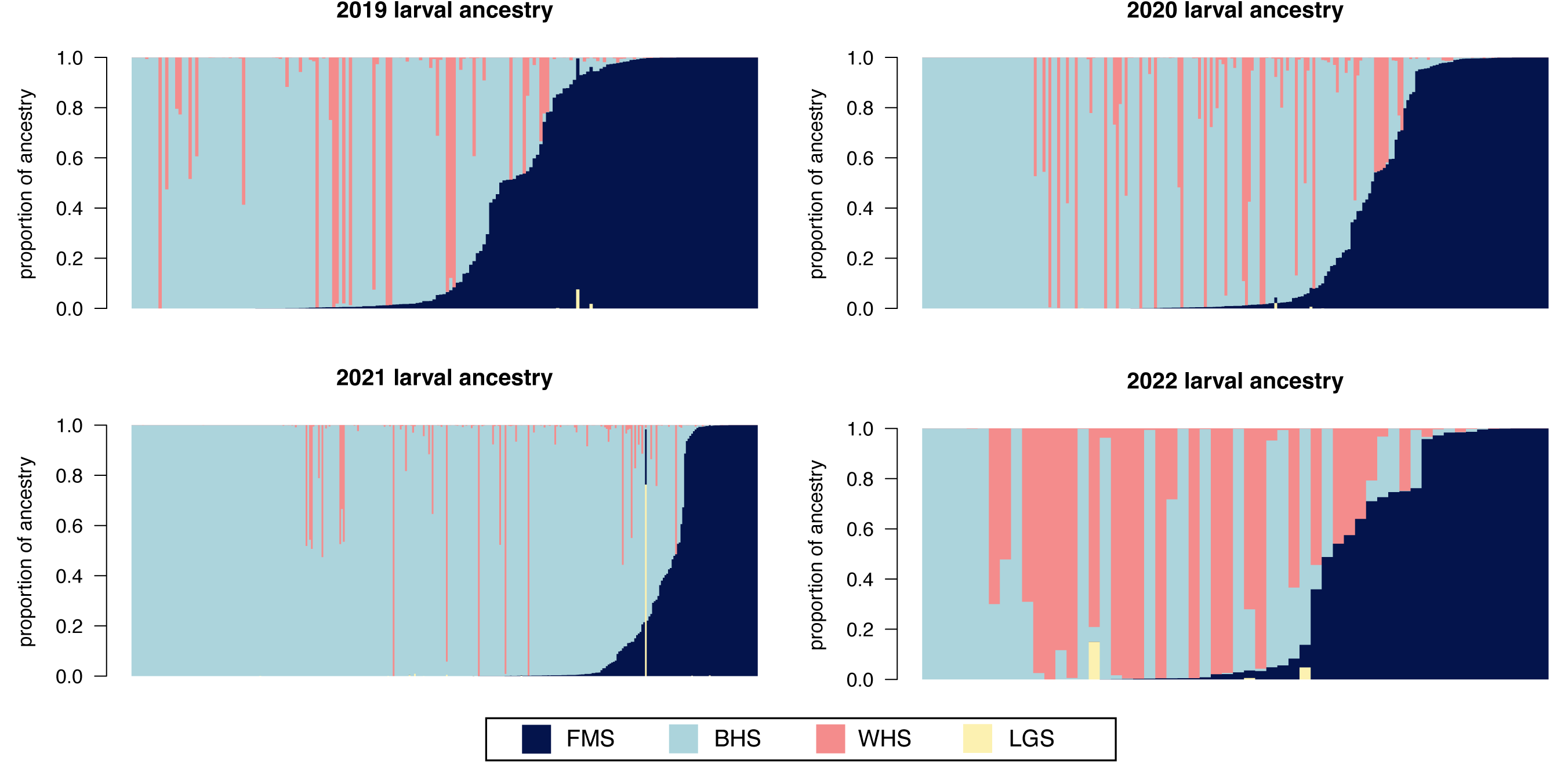
Larval ancestry from each of the four parental species across the four study years. Colours represent the four parental species, flannelmouth sucker (FMS), bluehead sucker (BHS), white sucker (WHS), and longnose sucker (LGS). Each bar along the x-axis represents an individual with their proportion of ancestry from the parental species along the y-axis. Sample size in 2019 is n=187, 2020 is n=213, 2021 is n=352, and 2022 is n=56.

### Variation in hybridization dynamics between species pairs

Combining estimates of *q* and *Q* allowed for a detailed analysis of hybridization between species pairs. With these estimates we could better understand which types of hybrids were present in the sampled population (F1 hybrids, F2 hybrids, backcrosses, etc.). Using the three species of greatest interest, flannelmouth, bluehead and white suckers, we evaluated hybridization between all possible species crosses.

Evaluation of white *×* flannelmouth suckers showed multiple F1 hybrids but no F2 hybrids (Fig. 4A). Additionally, backcrosses were seen more with hybrid individuals crossing back to the native flannelmouth sucker. 95% CIs (credible intervals) surrounding these data were extremely narrow with the mean CI for *q* being 0.004 and 0.0083 for *Q*.

**Figure 4:**
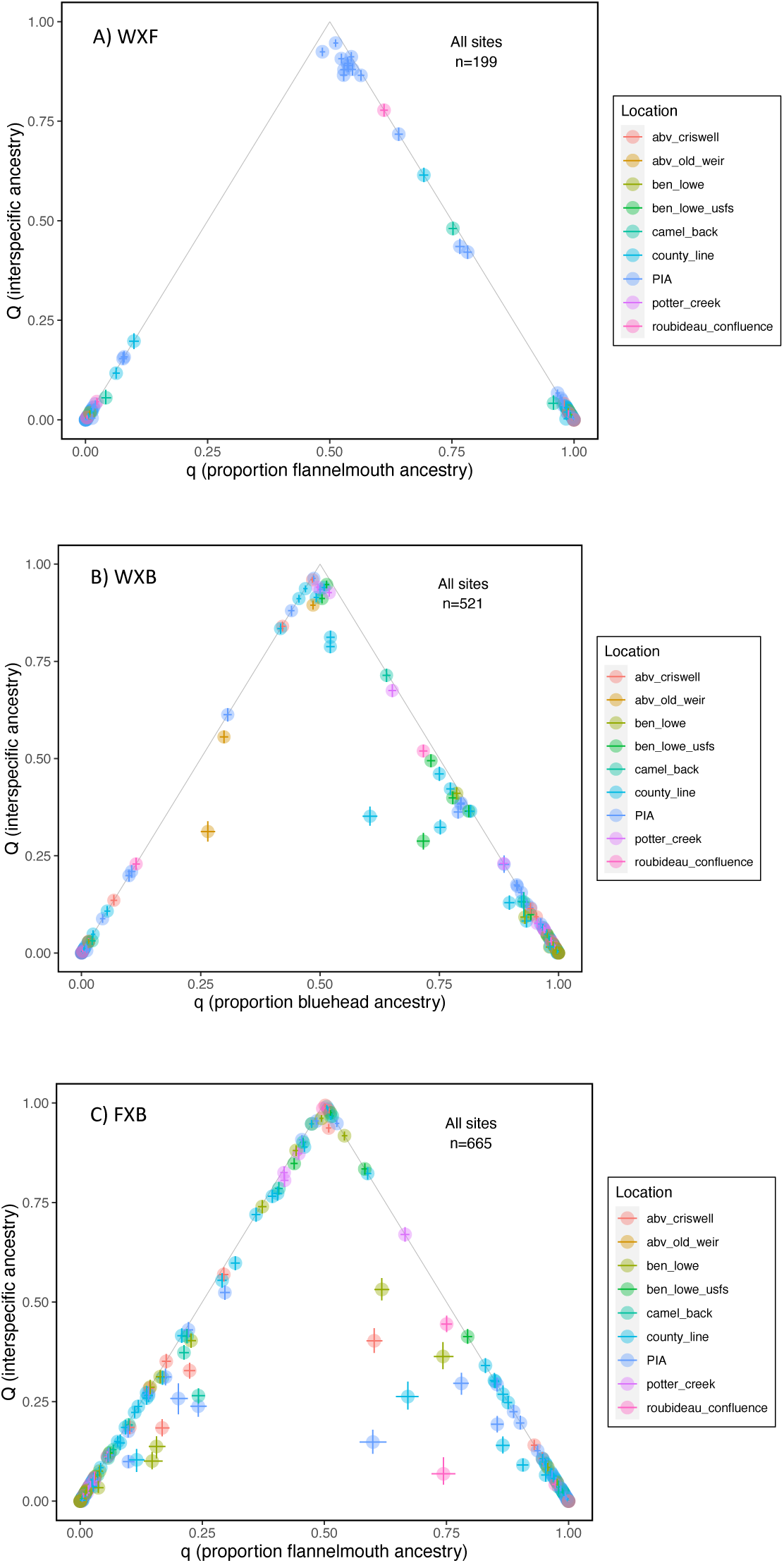
Estimates of *q* versus *Q* for the different hybrid crosses. Values of *q* represent the proportion of an individuals ancestry that comes from each parental species. *Q* represents interspecific ancestry, the proportion of loci in an individual that have ancestry from both parental species. A) Shows individuals with white or flannelmouth ancestry, B) shows individuals with white or bluehead ancestry, and C) shows individuals with flannelmouth or bluehead ancestry. F1 individuals can be seen at the top of the triangle (*q* =0.5, *Q* =1). F2 individuals are in the middle (*q* =0.5, *Q* =0.5) and backcrosses are along the triangle lines (*q* =0.25 or 0.75, *Q* =0.5).

Analysis of white *×* bluehead sucker crosses showed many hybrids being F1 hybrids or backcrosses (Fig. 4B). Backcrossed individuals were seen towards both white and bluehead suckers, although more hybrids were backcrossing with the native bluehead sucker. Narrow 95% CIs were observed with *q* and *Q* having mean CIs of 0.0035 and 0.0067, respectively.

Flannelmouth *×* bluehead sucker crosses were also evaluated. We saw many F1 and backcrossed individuals between this species pair (Fig. 4C). Backcrosses appear to go in both directions with hybrids crossing back to either of the native flannelmouth or bluehead suckers. Again, 95% CIs remained narrow with means of 0.0049 for *q* and 0.0090 for *Q*.

### Adult identification in the field

To account for error in identification by field personnel, a reference and a random subset of adult fish moving upstream to spawn were sampled for genetic analysis. A phenotypic identification was recorded and a fin clip was taken for a genetic identification comparison. By getting estimates of human error in identification we can account for this error when looking at how well the weir worked.

Phenotypic identification of adult fish varied by species. The frequencies of pure species (flannelmouth, bluehead, and white sucker) were over-represented in the field identification data, while the frequencies of hybrids were under-represented. Combining both the randomly collected adult fish and the reference adult fish, white suckers were identified 49 times phenotypically but only 34 times genetically. Bluehead suckers were identified 68 times phenotypically but 56 times genetically. Flannelmouth suckers were identified 80 times in the field and only 56 times genetically. Hybrids between white *×* bluehead suckers were identified 2 times phenotypically but 21 times using genetics. Hybrids between white *×* flannelmouth suckers were identified 2 times in the field but 7 times genetically. Flannelmouth *×* bluehead suckers, longnose *×* flannelmouth suckers, longnose *×* bluehead suckers, and other multi-species hybrid crosses were identified 0 times phenotypically in the field. However, genetic identification found 16, 1, 2, and 7 of these hybrids, respectively (Table 1).

**Table 1:** Summary of field vs genetic identification given to adult *Catostomus* fish. White *×* bluehead sucker hybrid (WXB), white *×* flannelmouth sucker hybrid (WXF), flannelmouth *×* bluehead hybrid (FXB), longnose *×* flannelmouth sucker hybrid (LXF), and longnose *×* bluehead sucker hybrid (LXB).

| Ancestry | Field ID<br>(Random) | Genetic ID<br>(Random) | Field ID<br>(Reference) | Genetic ID<br>(Reference) | Field ID<br>(All) | Genetic ID<br>(All) |
| --- | --- | --- | --- | --- | --- | --- |
| White | 1 | 1 | 48 | 33 | 49 | 34 |
| Bluehead | 30 | 25 | 38 | 33 | 68 | 56 |
| Flannelmouth | 38 | 31 | 42 | 25 | 80 | 56 |
| Longnose | 0 | 0 | 9 | 5 | 9 | 5 |
| WXB | 2 | 4 | 0 | 9 | 2 | 21 |
| WXF | 2 | 4 | 0 | 11 | 2 | 7 |
| FXB | 0 | 4 | 0 | 12 | 0 | 16 |
| LXF | 0 | 0 | 0 | 1 | 0 | 1 |
| LXB | 0 | 0 | 0 | 2 | 0 | 2 |
| Other | 0 | 4 | 0 | 3 | 0 | 7 |

## Discussion

### Weir success

Overall, the weir had no statistically significant effect on mean proportion of white sucker ancestry across varying degrees of weir usage (Fig. 2). However, when the weir was present for the entire spawning season we saw the lowest amount of white sucker ancestry compared to any other year at 3.59% (Fig. 1, Table S1). This is a positive result and shows that when fully active, the weir appears to be successful at excluding non-native suckers. However, in 2020 and 2022 when the weir had to be pulled from the creek early, white sucker ancestry was 11.75% and 37.80%, respectively. This suggests that if the weir was not present for the entire spawning season, its effect was variable and in 2022 specifically, the weir’s effect was negligible. This information portrays the importance of using the weir for the entire spawning season. If the weir cannot withstand an entire season in the creek, then it is likely not going to have a significant effect on reducing non-native fish. Snowfall and runoff conditions in the Western U.S. are highly variable, and are predicted to become more-so with climate change. For instance, between 2018 and 2023, both record low and high snowpack occurred in the Roubideau Creek drainage. Likewise, hotter and cooler than usual springs have occurred within the same period greatly altering how fast and heavy runoff has occurred. These conditions are difficult to predict, which makes planning and implementing a conservation intervention on the scale of the RBW difficult if it is likely to fail under higher than average flow conditions. Our finding that partial implementation had no detectable effect does detract from the attractiveness of this particular device as a conservation action. However, the fact that less than 4% ancestry was attributed to white sucker in the full implementation year (2021) does show promise that a more robust design could effectively limit hybridization in the system.

Across all years, bluehead suckers, followed by flannelmouth suckers, made up the highest proportion of larval ancestry, except for in 2022 when white suckers dominated (Fig. 3). Sample size between years varied substantially from a high of 352 individuals in 2021, to a low of 56 individuals in 2022 (due to sample preservation issues in the field). It is possible that more consistent sample size across years could produce slightly different results. However, these results do show hybridization between both native fish, flannelmouth and bluehead sucker, with the non-native white sucker across all years (Fig. 3).

We would expect that the removal of white suckers in each year would reduce the population of potential white sucker spawners in Roubideau Creek the following year. A high proportion of suckers appear to spawn annually and at the same site (Hooley-Underwood et al. 2019); therefore, one would expect the weir to have a continued effect on the following year. However, we did not observe this compounding annual effect. In 2021, the weir was active for the entire spawning season, and we saw a very low proportion of non-native ancestry in larvae that year. However, the following year, when the weir was in for 48 days before being removed, had the most non-native ancestry within the sampled larval population of all four years. This might suggest the need for continued interventions yearly until the non-native species are no longer a threat to the native fishes.

Our control year, 2019, also provides an area for more exploration. In 2019, the weir was not present, yet white sucker ancestry in the sampled population was lower than 2020 and 2022 when the weir was in the creek for part of the season. Baseline white sucker ancestry was less than 10% of the entire population in the control year. This might suggest that another external factor not accounted for is impacting which fish are using the spawning habitat every year. It is unknown to what extent hybridization dynamics vary across years in the absence of an intervention.

During periods of weir operation, the majority of adult suckers captured appeared phenotypically to be of native ancestry (84% in 2020, 90.8% in 2021, 90.4% in 2022). While 2020 and 2022 represent incomplete efforts, our results do suggest that the ratio of field-identified parental species reflected larval ancestry in 2020 and 2021, but not 2022. The previously mentioned sample size issue may explain some of the disconnect in 2022, but this disconnect may further point to external factors that favor the production of non-native ancestry under certain unidentified conditions.

### Hybridization between species pairs

Hybridization dynamics between the three most abundant species pairs varied. We saw no indication of F2 hybrids between flannelmouth and white suckers, in contrast to what has been previously seen (Mandeville et al. 2017). Additionally, white *×* flannelmouth backcrossed individuals appeared to be backcrossing in the direction of the native flannelmouth suckers. White *×* bluehead crosses also showed this same trend, with backcrossed individuals leaning towards the native bluehead sucker. This asymmetrical backcrossing was not identified in previous work in these species (Mandeville et al. 2017) and begs the question, is there something specific to this system causing backcrossing to lean towards the native fish? We would expect that the exclusion of white suckers and hybrids from upstream spawning habitat alone might explain this; however, when looking at data from only 2019 (no intervention present) we still appear to see backcrossing leaning to the native fish (Fig. S4, Fig. S5). However, sample size and amount of backcrossed individuals were low in 2019 alone; therefore, more data will be necessary to understand this trend better. When looking at the native cross, flannelmouth *×* bluehead suckers, we see backcrossing in both directions. Additionally, we identified multiple F1 individuals.

Another trend that is consistent across all species pairs is the relative lack of F2 hybrids. Compared to F1 and backcrossed hybrids, there are notably fewer F2 individuals. One potential explanation for this observation is hybrid breakdown. Hybrid breakdown is a phenomenon where F1 hybrids are viable and fertile, however, F2 or later generation hybrids have reduced fitness or are inviable. This would explain the lack of F2 hybrids seen in this study. Hybrid breakdown has been documented in other fish species like cichlids (Stelkens et al. 2015), trout (Muhlfeld et al. 2009), and swordtail fishes (Payne et al. 2022), making this an interesting line for further investigation.

### Flow

A key difference between years that needs to be explored further is flow conditions. This river system, and the surrounding systems that this study was located in are very hydrologically flashy and experience rapid changes in flow conditions. Exploring the impact flow has on different species of suckers might provide an opportunity to understand if flow had a significant effect on which species were present across the four study years, or if different levels of flow alter reproductive outcomes of different parental species, and therefore hybridization rates. If flow is an important factor in sucker population composition, it may be interesting to look further into the swimming abilities of the species of interest. Currently, our understanding is that the critical swimming velocities of native flannelmouth and bluehead suckers are significantly different than that of the non-native white sucker (Underwood et al. 2014). Additionally, when under constant acceleration conditions, bluehead suckers had the best swimming ability, with flannelmouth suckers having the worst, and white suckers being intermediate (Underwood et al. 2014). Interestingly, near record levels of runoff occurred in both 2019 and 2022, but we observed low white sucker ancestry in 2019 and high white sucker ancestry in 2022. One difference of note is that spring warming was relatively typical in 2019, but abnormally heavy snowpack drove the heavy runoff, while in 2022, snowpack was near average, but spring warming was delayed followed by an extremely rapid warming period that led to a late but heavy runoff. In contrast, 2020 and 2021 were both sub-average snowpack and runoff years, and we saw a relatively large difference in native sucker ancestry produced in those two years. The complexities of these hydrographical dynamics are difficult to detangle in our study system, but likely affected our results, as well as reproductive outcomes of suckers in the system. Going forward, investigating interconnectivity between the hydrograph and ancestry of the larval population might provide interesting insights into the trends seen in this study.

### Adult fish identification

Phenotypic identification of adult fish was less accurate than expected. Incorrect identifications came in all forms. Many times hybrids were being identified as one of the pure parental species, and therefore, numbers of hybrids being identified were extremely low. This supports the idea that phenotypically identifying hybrids can be difficult, especially when these hybrids are advanced generation or backcrosses (Thompson et al. 2021). In this study, we did see that generally, more advanced generation hybrids were identified as pure individuals, but several F1 hybrids were also misidentified. Often hybrids can be cryptic and express phenotypes that resemble one parental species more (Thompson et al. 2021), making it increasingly difficult to ID these fish. However, we also saw pure adult species being identified incorrectly as another pure species, further showing the importance of genetic evaluation for species identification. We do suspect that some misidentifications were due to misspeak or data transcription errors associated with the fast pace necessary to process at times thousands of fish in under a 12 hour period. This represents another challenge of operating a conservation intervention on this scale where both inconsistent phenotypic representation of genotype and human error are likely to result in the passage of some non-native and hybridized adults. Overall, a total of 26.67% of adult fish were incorrectly identified in the field (Table S2). Additionally, entropy plots of adult fish in 2021 and 2022 can be seen to visualize composition of sampled adults (Fig. S6) and a supplementary table can be used to look at field versus genetic ID of each individual fish (Table S2). One notable feature of the adult data is that the proportions of adult fish varies substantially across years.

### Conclusions and future directions

The RBW used in this study did not have any significant effect on proportion of white sucker ancestry within the sampled larval population across varying degrees of weir usage (*p >* 0.05, R=0). Ancestry of the sampled population varied across years with the amount of ancestry coming from each of the four parental species fluctuating across all four years. The unpredictability of the hydrograph in this river system plays a huge role in the ability of the weir to be operable for an entire season. Going forward, further research could be focused on finding an intervention that is better suited to intermittent streams like Roubideau Creek, which can experience flashy flow changes and problematic debris loads.

One important takeaway from this research is that this study demonstrates the importance of multiyear studies. If we only had sampled one year with the resistance board weir intervention and one without, the results from this study might look a lot different. For example, if only 2021 (full weir year) and 2022 (partial weir year) were sampled, the change in proportion of white sucker ancestry within the sampled population would appear to be drastically different (3.59% to 37.80%). In this case, one would conclude that the weir had a significant effect on proportion of white sucker ancestry. However, when we look at more years, like 2019 and 2020, we can see that proportion of white sucker ancestry fluctuates across years for unknown reasons. Having a study that spans multiple years allowed us to see this information that would potentially have been missed if the study was shorter.

As mentioned, clearly there is another factor influencing white sucker ancestry as the proportions vary widely across years. For example, the two partially controlled years had 11.75% and 37.80% white sucker ancestry in the sampled population, with the higher proportion coming from the year that was partially controlled for a longer period of time - the opposite of what we might expect if length of weir operation drives reduction in white sucker ancestry in larval fish. Additional study would be needed to identify what is causing these changes in population composition between years.

Another line of research continuing from this project would be considering the additive effect of one year on another. It would be interesting to see if multiple years of weir use have an additive effect on one another, therefore causing increasingly lower percentages of white sucker ancestry within the sampled population as years go by. Movement data collected from PIT tagged fish within this drainage show that individuals of both native and non-native sucker species are likely to spawn within the same tributary every year that environmental conditions allow (Hooley-Underwood et al. 2019). Therefore, we would expect an additive effect over multiple years of use. Currently, our findings show that the year where the weir was present for the entire spawning season had the lowest amount of white sucker ancestry in the population. However, the following year where the RBW was active for only part of the season, had the highest amount of white sucker ancestry. This would suggest that one year does not have a strong effect on the following year. However, more years of sustained weir use is necessary to draw conclusions on this topic.

This study also demonstrated that field identification of adult suckers is challenging, especially with hybrids involved. Pure species were consistently over identified in the field and hybrids under identified in the field. Morphological traits that field staff in the region rely on such as mouth structure, body shape, scale counts and organization, and fin shape and ray counts appear to be less definitive than they have been previously thought. Many of the meristic based diagnostics are time consuming. Our study demonstrates that having high identification accuracy may be extremely difficult when handling large numbers of catostomids with limited time. If individuals in the field are better able to identify hybrids, especially cryptic-looking hybrids, the success of the weir might be increased. Even so, hybrids can have intermediate traits in things like head depth, which relies on specific measurements, morphological identification may always be problematic (Thompson et al. 2021).

To conclude, a resistance board weir as a conservation intervention in this river system did not provide any significant effect on the removal of non-native sucker fish. However, we did see the lowest proportion of white sucker ancestry when the weir was present for the entire spawning season, suggesting the potential for it to be successful if implementation could be more consistent. The limiting factor for the success of the weir is its ability to withstand high flow conditions. Future efforts should focus on alternative exclusion devices and understanding to what extent sustained exclusion may result in an additive effect.

## Data accessibility

With acceptance of this manuscript, all data and scripts will be publicly available on Data Dryad, NCBI SRA, and GitHub (https://github.com/jcampb37).

## Author contributions

JNC and EGM planned and designed research analyses, created genomic data, and performed bioinformatic processes. ZEHU participated in the project conceptualization and study design, and oversaw field sampling and weir operations. All authors contributed to writing and revising of the manuscript.

## Conflict of Interest

The authors have no conflict of interest to declare.

## Acknowledgements

This work was funded by Colorado Parks and Wildlife (Species Conservation Trust Fund Project Number: SCA20B). Computing for this project was successfully completed using a RRG Allocation on the Digital Research Alliance of Canada’s Cedar cluster to E.G. Mandeville. We sincerely thank Kevin Thompson for his extensive work in the drainage that led to this project, and for conceptualizing, initiating, and participating in this whole endeavor. We thank numerous Colorado Parks and Wildlife, Bureau of Land Management, and U.S. Fish and Wildlife Service biologists and seasonal technicians, as well as several outstanding volunteers for doing the hard work of operating the weir and collecting all genetic samples. This manuscript was improved by comments from members of the Mandeville Lab at University of Guelph, as well as Eryn McFarlane, Rob McLaughlin, and Sally Adamowicz.

---

**Figure S1:**
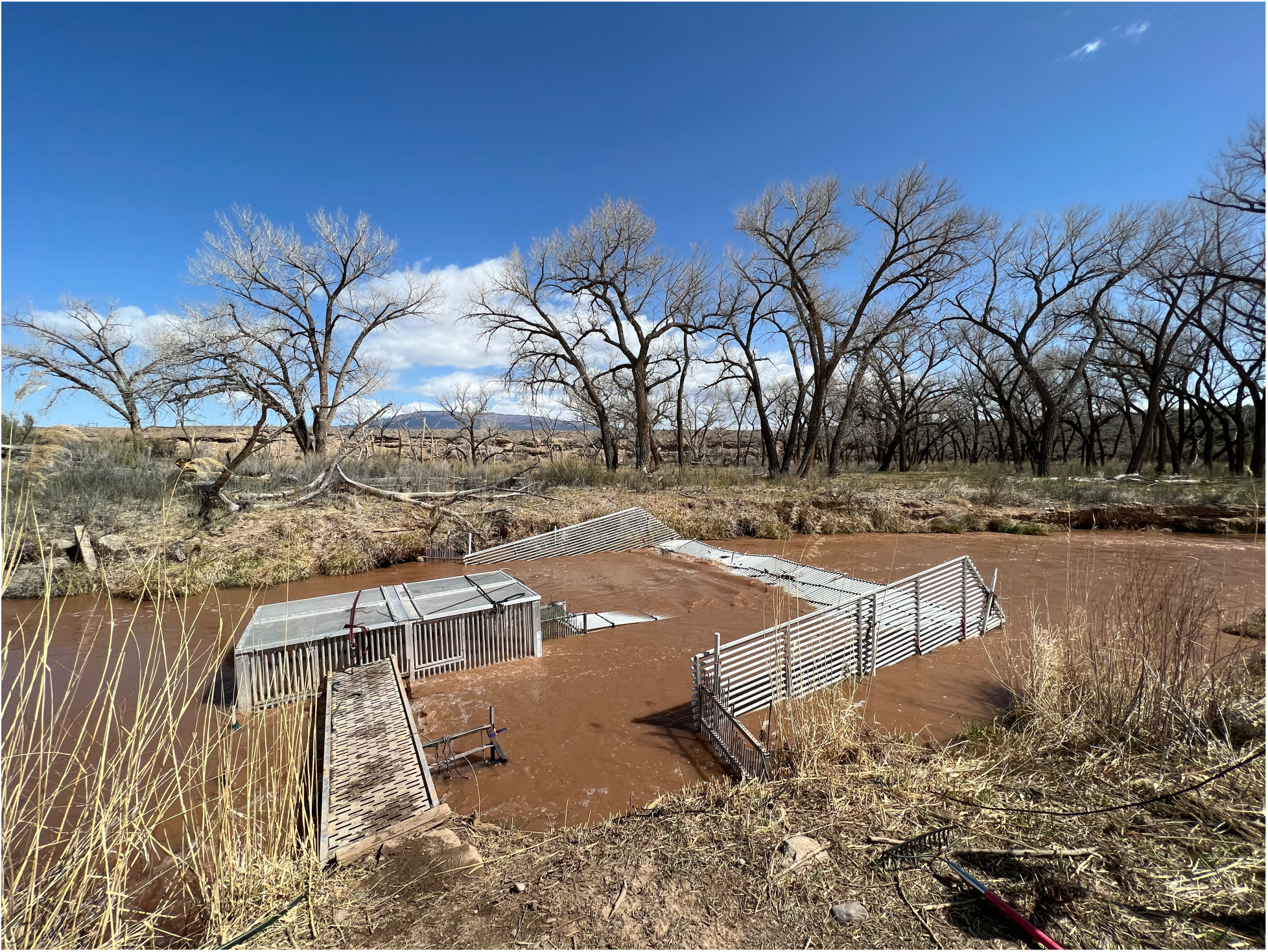
Photo of the resistance board weir installed in Roubideau Creek during 2022. Fish swim into the weir from the right of the image and are funneled into the cage in the center. Field technicians remove fish from the cage, ID them, then determine if the fish are allowed to pass into spawning habitat (upstream of the left side of the photo).

**Figure S2:**
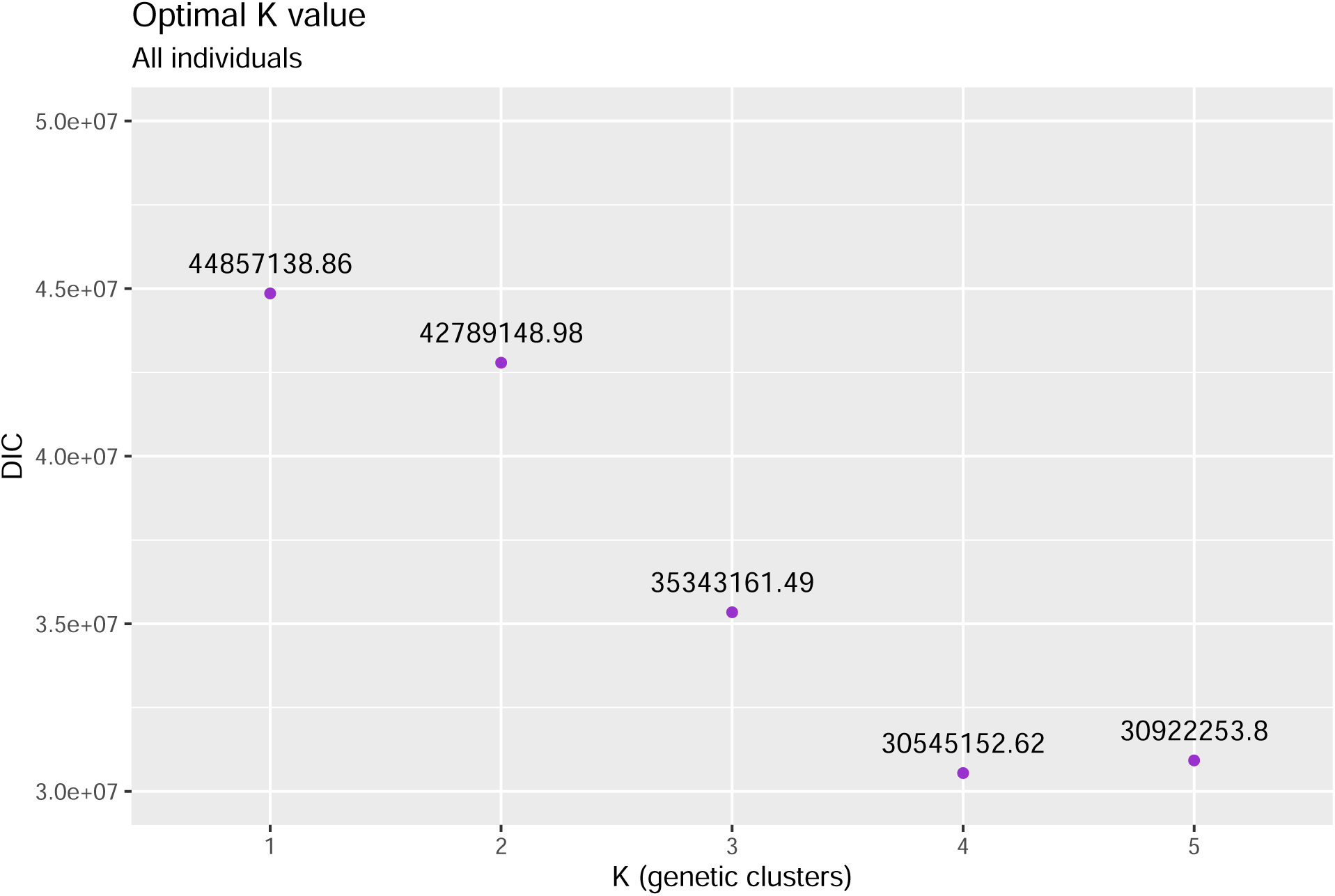
Optimal *k* value for runs of entropy as determined by DIC (Deviance Information Criterion) values. The *k* value with the lowest DIC value is the *k* value for which the model has determined is the best fit. Here *k* = 4 is most supported.

**Figure S3:**
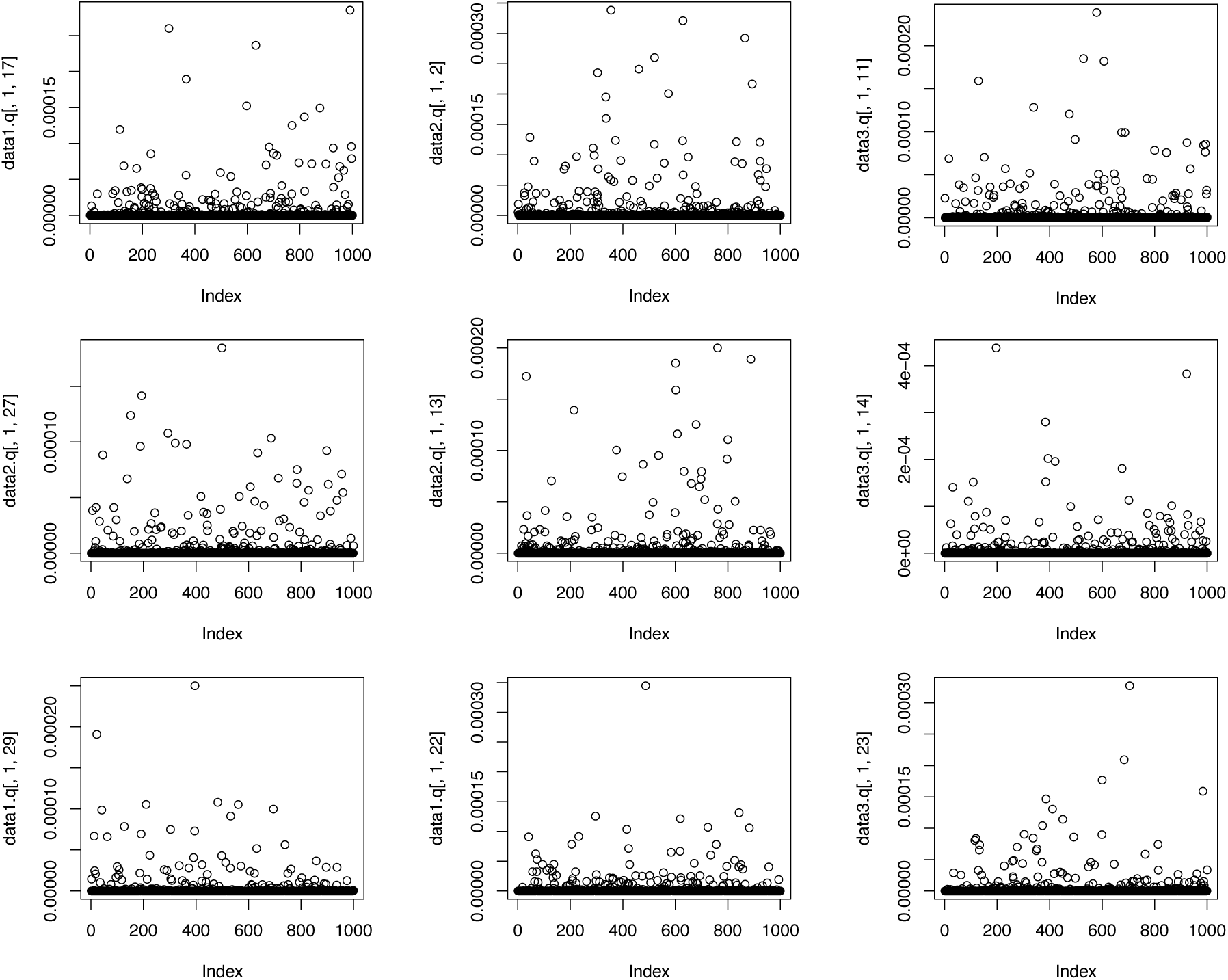
Random trace plots from MCMC (Markov chain Monte Carlo) chains demonstrating proper model convergence by entropy.

**Figure S4:**
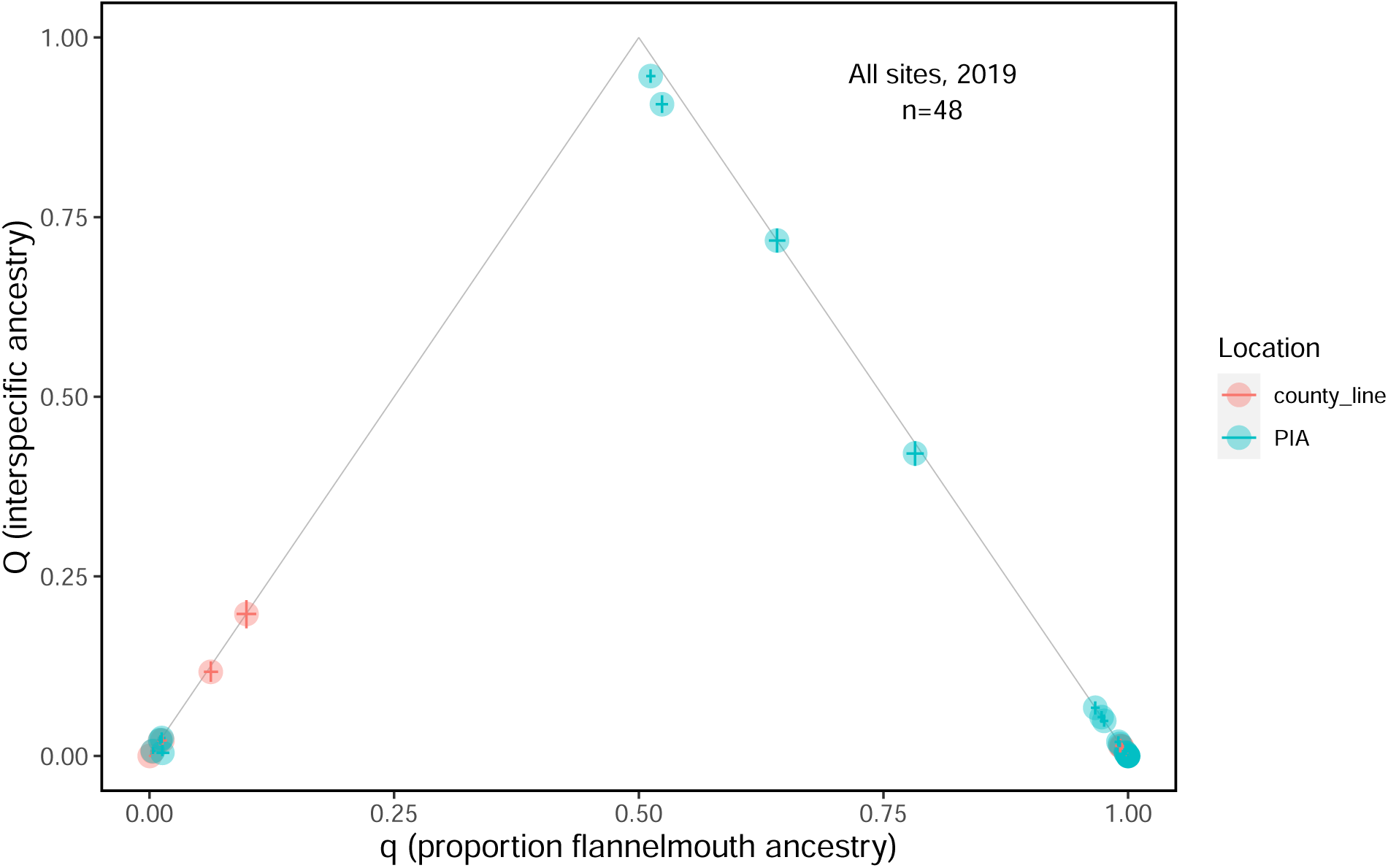
White*×*flannelmouth sucker estimates of *q* versus *Q* for 2019. F1 individuals can be seen at the top of the triangle (*q* =0.5, *Q* =1). F2 individuals are in the middle (*q* =0.5, *Q* =0.5) and backcrosses are along the triangle lines (*q* =0.25 or 0.75, *Q* =0.5).

**Figure S5:**
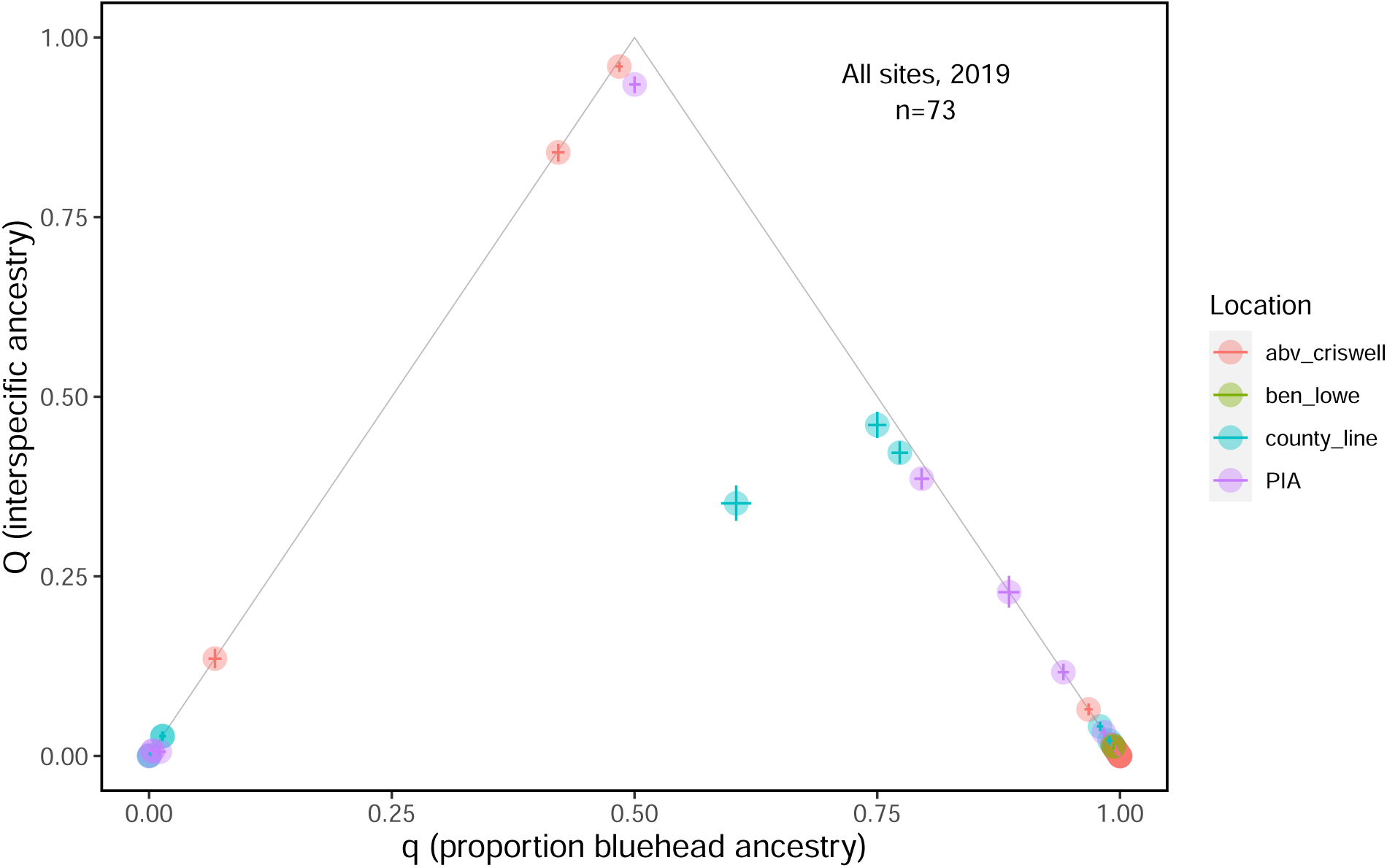
White*×*bluehead sucker estimates of *q* versus *Q* for 2019. F1 individuals can be seen at the top of the triangle (*q* =0.5, *Q* =1). F2 individuals are in the middle (*q* =0.5, *Q* =0.5) and backcrosses are along the triangle lines (*q* =0.25 or 0.75, *Q* =0.5).

**Figure S6:**
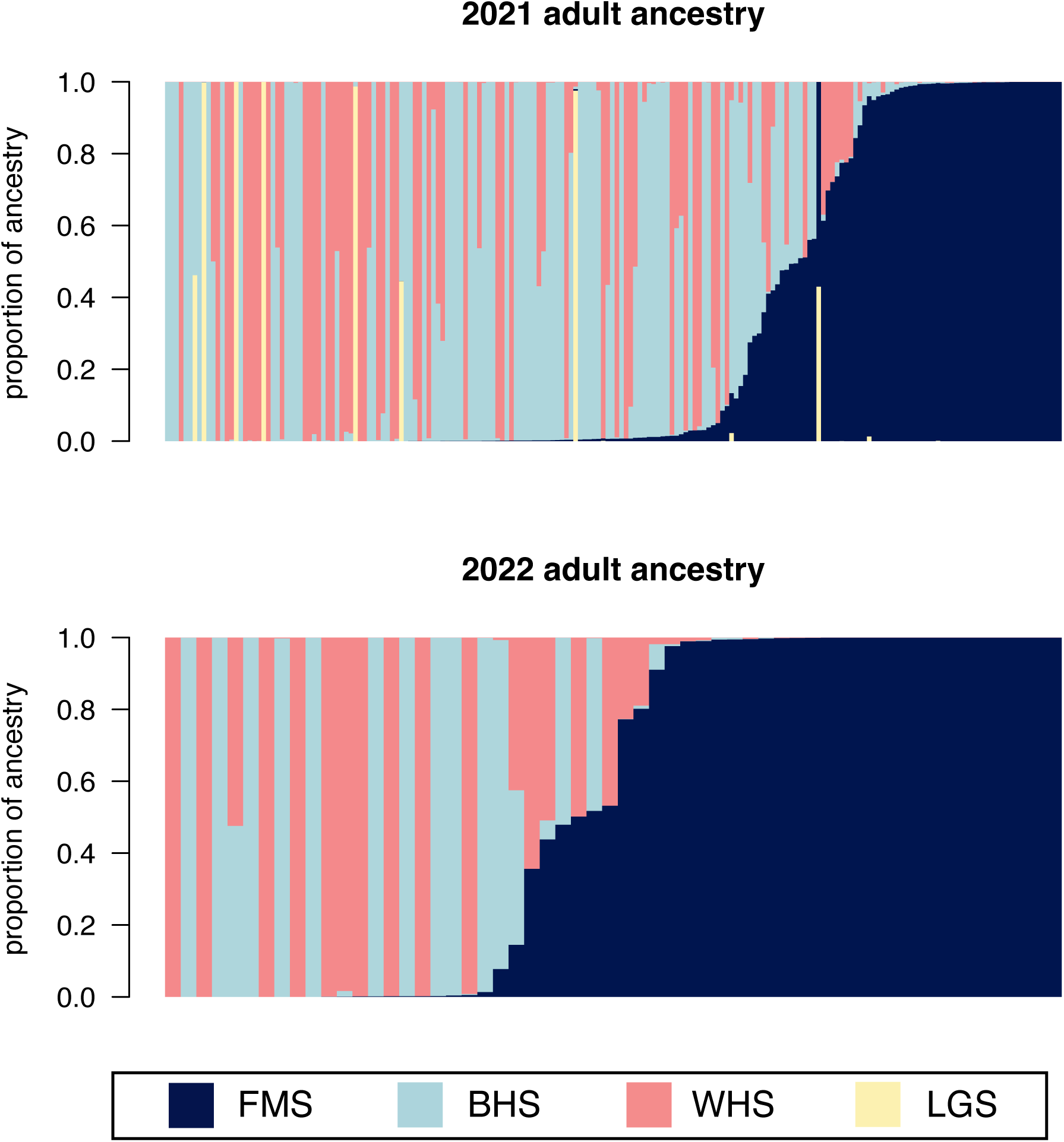
Ancestry estimates of parental species by entropy. Colours represent the four parental species, flannelmouth sucker (FMS), bluehead sucker (BHS), white sucker (WHS), and longnose sucker (LGS). Each line along the x-axis represents an individual with their proportion of ancestry from the parental species along the y-axis. In 2021, some additional individuals are included relative to Table 1; these individuals were juveniles so were not included in our estimates of phenotypic identification success.

**Table S1:**
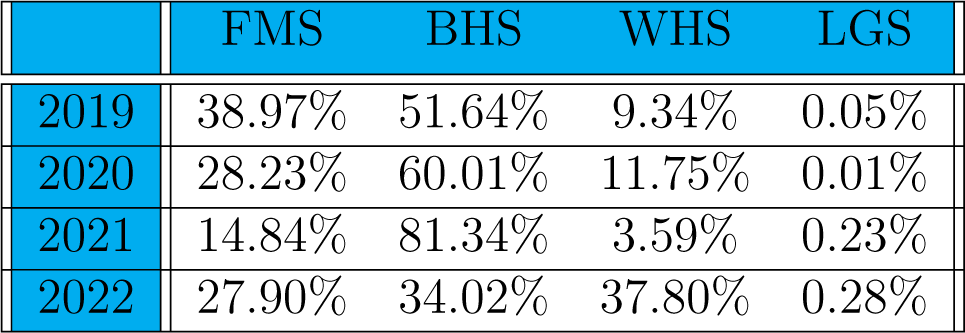
Percentage of sampled larval population with ancestry from each parental species across years. Flannelmouth sucker (FMS), bluehead sucker (BHS), white sucker (WHS), and longnose sucker (LGS).

|  | FMS | BHS | WHS | LGS |
| --- | --- | --- | --- | --- |
| 2019 | 38.97% | 51.64% | 9.34% | 0.05% |
| 2020 | 28.23% | 60.01% | 11.75% | 0.01% |
| 2021 | 14.84% | 81.34% | 3.59% | 0.23% |
| 2022 | 27.90% | 34.02% | 37.80% | 0.28% |

**Table S2:** Supplemental table showing the field phenotypic ID, genetic ID, and proportional ancestry (*q* from entropy) for each adult sucker sampled. Flannelmouth sucker (FMS), bluehead sucker (BHS), white sucker (WHS), white*×*flannelmouth sucker (WXF), white*×*bluehead sucker (WXB), and flannelmouth*×*bluehead sucker (FXB).

| Individual | Bluehead | White | Longnose | Flannemouth | Genetic | Field |
| --- | --- | --- | --- | --- | --- | --- |
| 210323BHS03 | 0.91 | 0.00 | 0.00 | 0.09 | FXB | BHS |
| 210323BHS04 | 1.00 | 0.00 | 0.00 | 0.00 | BHS | BHS |
| 210323BHS05 | 0.00 | 0.00 | 0.00 | 1.00 | FMS | BHS |
| 210323BHS06 | 0.98 | 0.00 | 0.00 | 0.01 | BHS | BHS |
| 210323BHS07 | 0.50 | 0.50 | 0.00 | 0.00 | WXB | BHS |
| 210323BHS08 | 0.81 | 0.00 | 0.00 | 0.18 | FXB | BHS |
| 210323BHS09 | 1.00 | 0.00 | 0.00 | 0.00 | BHS | BHS |
| 210323BHS10 | 1.00 | 0.00 | 0.00 | 0.00 | BHS | BHS |
| 210323BHS11 | 0.98 | 0.01 | 0.00 | 0.01 | BHS | BHS |
| 210323BHS12 | 1.00 | 0.00 | 0.00 | 0.00 | BHS | BHS |
| 210323BHS13 | 0.97 | 0.02 | 0.00 | 0.01 | BHS | BHS |
| 210323BHS14 | 1.00 | 0.00 | 0.00 | 0.00 | BHS | BHS |
| 210323BHS15 | 0.97 | 0.00 | 0.00 | 0.03 | BHS | BHS |
| 210323BHS16 | 0.93 | 0.06 | 0.00 | 0.01 | WXB | BHS |
| 210323BHS17 | 1.00 | 0.00 | 0.00 | 0.00 | BHS | BHS |
| 210323BHS18 | 1.00 | 0.00 | 0.00 | 0.00 | BHS | BHS |
| 210323BHS19 | 1.00 | 0.00 | 0.00 | 0.00 | BHS | BHS |
| 210323BHS20 | 1.00 | 0.00 | 0.00 | 0.00 | BHS | BHS |
| 210323BHS21 | 1.00 | 0.00 | 0.00 | 0.00 | BHS | BHS |
| 210323BHS22 | 1.00 | 0.00 | 0.00 | 0.00 | BHS | BHS |
| 210323BHS23 | 0.99 | 0.00 | 0.00 | 0.01 | BHS | BHS |
| 210323BHS24 | 1.00 | 0.00 | 0.00 | 0.00 | BHS | BHS |
| 210323BHS25 | 1.00 | 0.00 | 0.00 | 0.00 | BHS | BHS |
| 210323BHS26 | 1.00 | 0.00 | 0.00 | 0.00 | BHS | BHS |
| 210323BHS27 | 1.00 | 0.00 | 0.00 | 0.00 | BHS | BHS |
| 210323BHS28 | 0.97 | 0.00 | 0.00 | 0.03 | BHS | BHS |
| 210323BHS29 | 1.00 | 0.00 | 0.00 | 0.00 | BHS | BHS |
| 210323BHS30 | 1.00 | 0.00 | 0.00 | 0.00 | BHS | BHS |
| 210323FMS01 | 0.07 | 0.05 | 0.00 | 0.88 | Other Hybrid | FMS |
| 210323FMS02 | 0.88 | 0.00 | 0.00 | 0.12 | FXB | FMS |
| 210323FMS03 | 0.56 | 0.00 | 0.00 | 0.44 | FXB | FMS |
| 210323FMS04 | 0.01 | 0.22 | 0.00 | 0.77 | WXF | FMS |
| 210323FMS05 | 0.02 | 0.37 | 0.00 | 0.61 | WXF | FMS |
| 210323FMS06 | 0.00 | 0.00 | 0.00 | 1.00 | FMS | FMS |
| 210323FMS07 | 0.00 | 0.21 | 0.00 | 0.79 | WXF | FMS |
| 210323FMS08 | 0.70 | 0.00 | 0.00 | 0.30 | FXB | FMS |
| 210323WHS01 | 0.02 | 0.98 | 0.00 | 0.00 | WHS | WHS |
| 210323WHS02 | 0.00 | 1.00 | 0.00 | 0.00 | WHS | WHS |
| 210323WHS03 | 0.92 | 0.08 | 0.00 | 0.00 | WXB | WHS |
| 210323WHS04 | 0.00 | 1.00 | 0.00 | 0.00 | WHS | WHS |
| 210323WHS05 | 0.01 | 0.99 | 0.00 | 0.00 | WHS | WHS |
| 210323WHS06 | 0.01 | 0.99 | 0.00 | 0.00 | WHS | WHS |
| 210323WHS07 | 0.54 | 0.46 | 0.00 | 0.00 | WXB | WHS |
| 210324FMS01 | 0.04 | 0.00 | 0.00 | 0.96 | FMS | FMS |
| 210324FMS02 | 0.00 | 0.22 | 0.00 | 0.78 | WXF | FMS |
| 210324FMS03 | 0.00 | 0.00 | 0.00 | 1.00 | FMS | FMS |
| 210324FMS04 | 0.00 | 0.00 | 0.00 | 1.00 | FMS | FMS |
| 210324FMS05 | 0.00 | 0.00 | 0.00 | 1.00 | FMS | FMS |
| 210324FMS06 | 0.52 | 0.00 | 0.00 | 0.48 | FXB | FMS |
| 210324FMS07 | 0.00 | 0.00 | 0.00 | 1.00 | FMS | FMS |
| 210324FMS08 | 0.00 | 0.00 | 0.00 | 1.00 | FMS | FMS |
| 210324FMS09 | 0.04 | 0.22 | 0.00 | 0.74 | WXF | FMS |
| 210324FMS10 | 0.01 | 0.03 | 0.00 | 0.96 | FMS | FMS |
| 210324FMS11 | 0.00 | 0.00 | 0.00 | 1.00 | FMS | FMS |
| 210324WHS01 | 0.00 | 0.99 | 0.00 | 0.00 | WHS | WHS |
| 210324WHS02 | 0.07 | 0.45 | 0.00 | 0.48 | Other Hybrid | WHS |
| 210324WHS03 | 0.00 | 0.99 | 0.00 | 0.00 | WHS | WHS |
| 210324WHS04 | 0.00 | 1.00 | 0.00 | 0.00 | WHS | WHS |
| 210326WHS01 | 0.00 | 1.00 | 0.00 | 0.00 | WHS | WHS |
| 210326WHS02 | 0.00 | 1.00 | 0.00 | 0.00 | WHS | WHS |
| 210327FMS01 | 0.01 | 0.00 | 0.00 | 0.99 | FMS | FMS |
| 210327FMS02 | 0.06 | 0.00 | 0.00 | 0.94 | FXB | FMS |
| 210327FMS03 | 0.03 | 0.00 | 0.00 | 0.97 | FMS | FMS |
| 210327FMS04 | 0.04 | 0.00 | 0.01 | 0.95 | WXF | FMS |
| 210327FMS05 | 0.16 | 0.00 | 0.00 | 0.84 | FXB | FMS |
| 210327FMS06 | 0.00 | 0.28 | 0.00 | 0.72 | WXF | FMS |
| 210327WHS01 | 0.00 | 0.99 | 0.00 | 0.01 | WHS | WHS |
| 210328FMS01 | 0.00 | 0.00 | 0.00 | 1.00 | FMS | FMS |
| 210328FMS02 | 0.00 | 0.00 | 0.00 | 1.00 | FMS | FMS |
| 210328FMS03 | 0.00 | 0.00 | 0.00 | 1.00 | FMS | FMS |
| 210328FMS04 | 0.44 | 0.00 | 0.00 | 0.56 | FXB | FMS |
| 210328FMS05 | 0.00 | 0.00 | 0.00 | 1.00 | FMS | FMS |
| 210328WHS01 | 0.54 | 0.46 | 0.00 | 0.00 | WXB | WHS |
| 210328WHS02 | 0.00 | 0.97 | 0.00 | 0.03 | WHS | WHS |
| 210328WHS03 | 0.00 | 1.00 | 0.00 | 0.00 | WHS | WHS |
| 210329WHS01 | 0.00 | 1.00 | 0.00 | 0.00 | WHS | WHS |
| 210329WHS02 | 0.61 | 0.37 | 0.00 | 0.02 | WXB | WHS |
| 210329WHS03 | 0.43 | 0.57 | 0.00 | 0.00 | WXB | WHS |
| 210329WHS04 | 0.01 | 0.99 | 0.00 | 0.00 | WHS | WHS |
| 210401WHS01 | 0.00 | 0.95 | 0.00 | 0.05 | WXF | WHS |
| 210402LGS01 | 0.55 | 0.00 | 0.45 | 0.00 | LXB | LGS |
| 210402WHS01 | 0.01 | 0.99 | 0.00 | 0.00 | WHS | WHS |
| 210402WHS02 | 0.00 | 1.00 | 0.00 | 0.00 | WHS | WHS |
| 210402WHS03 | 0.01 | 0.96 | 0.00 | 0.03 | WHS | WHS |
| 210402WHS04 | 0.00 | 0.99 | 0.00 | 0.01 | WHS | WHS |
| 210402WHS05 | 0.00 | 1.00 | 0.00 | 0.00 | WHS | WHS |
| 210402WHS06 | 0.00 | 1.00 | 0.00 | 0.00 | WHS | WHS |
| 210402WHS07 | 0.38 | 0.62 | 0.00 | 0.00 | WXB | WHS |
| 210402WHS08 | 0.00 | 1.00 | 0.00 | 0.00 | WHS | WHS |
| 210402WHS09 | 0.00 | 1.00 | 0.00 | 0.00 | WHS | WHS |
| 210403LGS01 | 0.00 | 0.00 | 1.00 | 0.00 | LGS | LGS |
| 210403LGS02 | 0.00 | 0.00 | 0.43 | 0.57 | LXF | LGS |
| 210407BHSR01 | 0.96 | 0.00 | 0.00 | 0.04 | BHS | BHS |
| 210407FMSR01 | 0.00 | 0.00 | 0.00 | 1.00 | FMS | FMS |
| 210407WHSR01 | 0.00 | 1.00 | 0.00 | 0.00 | WHS | WHS |
| 210407WXBR01 | 0.53 | 0.47 | 0.00 | 0.00 | WXB | WXB |
| 210408BHSR01 | 1.00 | 0.00 | 0.00 | 0.00 | BHS | BHS |
| 210408BHSR02 | 1.00 | 0.00 | 0.00 | 0.00 | BHS | BHS |
| 210408FMSR01 | 0.00 | 0.00 | 0.00 | 1.00 | FMS | FMS |
| 210408FMSR02 | 0.00 | 0.00 | 0.00 | 1.00 | FMS | FMS |
| 210409BHSR01 | 1.00 | 0.00 | 0.00 | 0.00 | BHS | BHS |
| 210409BHSR02 | 1.00 | 0.00 | 0.00 | 0.00 | BHS | BHS |
| 210409FMSR02 | 0.00 | 0.00 | 0.00 | 1.00 | FMS | FMS |
| 210410FMSR01 | 0.05 | 0.00 | 0.00 | 0.95 | WXF | FMS |
| 210410FMSR02 | 0.00 | 0.00 | 0.00 | 1.00 | FMS | FMS |
| 210411BHSR01 | 1.00 | 0.00 | 0.00 | 0.00 | BHS | BHS |
| 210411BHSR02 | 1.00 | 0.00 | 0.00 | 0.00 | BHS | BHS |
| 210411BHSR03 | 1.00 | 0.00 | 0.00 | 0.00 | BHS | BHS |
| 210411FMSR01 | 0.00 | 0.00 | 0.00 | 1.00 | FMS | FMS |
| 210412BHSR01 | 1.00 | 0.00 | 0.00 | 0.00 | BHS | BHS |
| 210412FMSR01 | 0.00 | 0.00 | 0.00 | 1.00 | FMS | FMS |
| 210412LGS01 | 0.00 | 0.00 | 1.00 | 0.00 | LGS | LGS |
| 210413BHSR01 | 1.00 | 0.00 | 0.00 | 0.00 | BHS | BHS |
| 210413FMSR01 | 0.00 | 0.00 | 0.00 | 1.00 | FMS | FMS |
| 210414BHSR01 | 0.43 | 0.57 | 0.00 | 0.01 | WXB | BHS |
| 210414BHSR02 | 1.00 | 0.00 | 0.00 | 0.00 | BHS | BHS |
| 210416BHSR01 | 1.00 | 0.00 | 0.00 | 0.00 | BHS | BHS |
| 210416BHSR02 | 1.00 | 0.00 | 0.00 | 0.00 | BHS | BHS |
| 210416FMSR02 | 0.01 | 0.00 | 0.00 | 0.99 | FMS | FMS |
| 210417BHSR01 | 0.99 | 0.00 | 0.00 | 0.01 | BHS | BHS |
| 210417BHSR02 | 1.00 | 0.00 | 0.00 | 0.00 | BHS | BHS |
| 210417BHSR03 | 1.00 | 0.00 | 0.00 | 0.00 | BHS | BHS |
| 210417BHSR04 | 1.00 | 0.00 | 0.00 | 0.00 | BHS | BHS |
| 210418BHSR01 | 1.00 | 0.00 | 0.00 | 0.00 | BHS | BHS |
| 210418BHSR02 | 1.00 | 0.00 | 0.00 | 0.00 | BHS | BHS |
| 210419FMSR01 | 0.50 | 0.00 | 0.00 | 0.50 | FXB | FMS |
| 210419FMSR02 | 0.00 | 0.00 | 0.00 | 1.00 | FMS | FMS |
| 210419LGS01 | 0.00 | 0.00 | 1.00 | 0.00 | LGS | LGS |
| 210420BHSR01 | 1.00 | 0.00 | 0.00 | 0.00 | BHS | BHS |
| 210420FMSR01 | 0.00 | 0.00 | 0.00 | 1.00 | FMS | FMS |
| 210420LGS01 | 0.49 | 0.00 | 0.00 | 0.51 | FXB | LGS |
| 210421BHSR01 | 0.82 | 0.05 | 0.02 | 0.11 | Other Hybrid | BHS |
| 210421FMSR01 | 0.00 | 0.00 | 0.00 | 1.00 | FMS | FMS |
| 210422BHSR01 | 1.00 | 0.00 | 0.00 | 0.00 | BHS | BHS |
| 210422FMSR01 | 0.71 | 0.00 | 0.00 | 0.29 | FXB | FMS |
| 210422FMSR02 | 0.00 | 0.30 | 0.00 | 0.70 | W XF | FMS |
| 210422FMSR03 | 0.00 | 0.00 | 0.00 | 1.00 | FMS | FMS |
| 210423BHSR01 | 1.00 | 0.00 | 0.00 | 0.00 | BHS | BHS |
| 210423BHSR03 | 1.00 | 0.00 | 0.00 | 0.00 | BHS | BHS |
| 210423FMSR01 | 0.00 | 0.00 | 0.00 | 1.00 | FMS | FMS |
| 210423LGS01 | 0.01 | 0.00 | 0.99 | 0.00 | LGS | LGS |
| 210424BHSR01 | 1.00 | 0.00 | 0.00 | 0.00 | BHS | BHS |
| 210424FMSR01 | 0.01 | 0.00 | 0.00 | 0.99 | FMS | FMS |
| 210424FMSR02 | 0.02 | 0.00 | 0.00 | 0.98 | FMS | FMS |
| 210424LGS01 | 0.54 | 0.00 | 0.46 | 0.00 | LXB | LGS |
| 210424WXBR01 | 0.53 | 0.47 | 0.00 | 0.00 | WXB | WXB |
| 210425BHSR01 | 0.79 | 0.06 | 0.00 | 0.15 | Other Hybrid | BHS |
| 210425BHSR02 | 0.44 | 0.28 | 0.00 | 0.28 | Other Hybrid | BHS |
| 210425BHSR03 | 0.54 | 0.46 | 0.00 | 0.00 | WXB | BHS |
| 210425FMSR01 | 0.00 | 0.00 | 0.00 | 1.00 | FMS | FMS |
| 210426BHSR01 | 0.99 | 0.00 | 0.00 | 0.01 | BHS | BHS |
| 210426FMSR01 | 0.00 | 0.00 | 0.00 | 0.99 | FMS | FMS |
| 210426LGS01 | 0.01 | 0.01 | 0.98 | 0.00 | LGS | LGS |
| BHS22032404 | 0.48 | 0.00 | 0.00 | 0.52 | FXB | BHS |
| BHS22032410 | 1.00 | 0.00 | 0.00 | 0.00 | BHS | BHS |
| BHS22032413 | 1.00 | 0.00 | 0.00 | 0.00 | BHS | BHS |
| BHS22032414 | 0.92 | 0.01 | 0.00 | 0.08 | FXB | BHS |
| BHS22032415 | 1.00 | 0.00 | 0.00 | 0.00 | BHS | BHS |
| BHS22032416 | 1.00 | 0.00 | 0.00 | 0.00 | BHS | BHS |
| BHS22032519 | 1.00 | 0.00 | 0.00 | 0.00 | BHS | BHS |
| BHS22032520 | 0.99 | 0.00 | 0.00 | 0.01 | BHS | BHS |
| BHS22032523 | 1.00 | 0.00 | 0.00 | 0.00 | BHS | BHS |
| BHS22032528 | 1.00 | 0.00 | 0.00 | 0.00 | BHS | BHS |
| FMS22032401 | 0.01 | 0.00 | 0.00 | 0.99 | FMS | FMS |
| FMS22032403 | 0.00 | 0.01 | 0.00 | 0.99 | FMS | FMS |
| FMS22032404 | 0.00 | 0.00 | 0.00 | 1.00 | FMS | FMS |
| FMS22032405 | 0.00 | 0.00 | 0.00 | 1.00 | FMS | FMS |
| FMS22032410 | 0.01 | 0.00 | 0.00 | 0.99 | FMS | FMS |
| FMS22032411 | 0.00 | 0.00 | 0.00 | 1.00 | FMS | FMS |
| FMS22032412 | 0.00 | 0.00 | 0.00 | 1.00 | FMS | FMS |
| FMS22032415 | 0.00 | 0.00 | 0.00 | 1.00 | FMS | FMS |
| FMS22032420 | 0.00 | 0.00 | 0.00 | 1.00 | FMS | FMS |
| FMS22032425 | 1.00 | 0.00 | 0.00 | 0.00 | BHS | FMS |
| FMS22032427 | 0.00 | 0.00 | 0.00 | 1.00 | FMS | FMS |
| FMS22032430 | 0.00 | 0.23 | 0.00 | 0.77 | W XF | FMS |
| FMS220405R02 | 0.00 | 0.00 | 0.00 | 1.00 | FMS | FMS |
| FMS220406R01 | 0.00 | 0.00 | 0.00 | 1.00 | FMS | FMS |
| FMS220406R03 | 0.00 | 0.00 | 0.00 | 1.00 | FMS | FMS |
| FMS220407R01 | 0.00 | 0.00 | 0.00 | 1.00 | FMS | FMS |
| FMS220408R01 | 0.00 | 0.00 | 0.00 | 1.00 | FMS | FMS |
| FMS220408R02 | 0.00 | 0.00 | 0.00 | 1.00 | FMS | FMS |
| FMS220409R01 | 0.01 | 0.19 | 0.00 | 0.80 | WXF | FMS |
| FMS220409R03 | 0.00 | 0.00 | 0.00 | 1.00 | FMS | FMS |
| FMS220409R04 | 0.07 | 0.02 | 0.00 | 0.91 | FXB | FMS |
| FMS220410R02 | 0.00 | 0.00 | 0.00 | 1.00 | FMS | FMS |
| FMS220411R02 | 0.00 | 0.00 | 0.00 | 1.00 | FMS | FMS |
| FMS220411R04 | 0.00 | 0.00 | 0.00 | 1.00 | FMS | FMS |
| FMS220416R01 | 0.00 | 0.00 | 0.00 | 1.00 | FMS | FMS |
| FMS220416R02 | 0.00 | 0.00 | 0.00 | 1.00 | FMS | FMS |
| FMS220417R02 | 0.01 | 0.02 | 0.00 | 0.98 | FMS | FMS |
| FMS220418R01 | 0.52 | 0.00 | 0.00 | 0.48 | FXB | FMS |
| WHS22032401 | 0.00 | 0.01 | 0.00 | 0.99 | FMS | WHS |
| WHS22032403 | 1.00 | 0.00 | 0.00 | 0.00 | BHS | WHS |
| WHS22032404 | 0.00 | 1.00 | 0.00 | 0.00 | WHS | WHS |
| WHS22032505 | 0.00 | 1.00 | 0.00 | 0.00 | WHS | WHS |
| WHS22032508 | 0.00 | 0.99 | 0.00 | 0.01 | WHS | WHS |
| WHS22032509 | 0.00 | 0.50 | 0.00 | 0.50 | WXF | WHS |
| WHS22032512 | 0.00 | 1.00 | 0.00 | 0.00 | WHS | WHS |
| WHS22032514 | 0.00 | 1.00 | 0.00 | 0.00 | WHS | WHS |
| WHS22032615 | 0.00 | 1.00 | 0.00 | 0.00 | WHS | WHS |
| WHS22032616 | 0.00 | 1.00 | 0.00 | 0.00 | WHS | WHS |
| WHS22032618 | 0.00 | 1.00 | 0.00 | 0.00 | WHS | WHS |
| WHS22032619 | 0.00 | 0.00 | 0.00 | 1.00 | FMS | WHS |
| WHS22032620 | 0.00 | 0.64 | 0.00 | 0.36 | WXF | WHS |
| WHS22032621 | 0.02 | 0.98 | 0.00 | 0.00 | WHS | WHS |
| WHS22032725 | 0.43 | 0.42 | 0.00 | 0.14 | Other Hybrid | WHS |
| WHS22032729 | 0.00 | 1.00 | 0.00 | 0.00 | WHS | WHS |
| WHS22032730 | 0.48 | 0.52 | 0.00 | 0.00 | WXB | WHS |
| WXF220408R02 | 0.05 | 0.51 | 0.00 | 0.44 | Other Hybrid | WXF |
| WXF220415R02 | 0.00 | 0.47 | 0.00 | 0.53 | WXF | WXF |

## Notes

### Competing Interest Statement

The authors have declared no competing interest.

